# Phage Peptides Mediate Precision Base Editing with Focused Targeting Window

**DOI:** 10.1101/2020.11.02.364430

**Authors:** Kun Jia, Yan-ru Cui, Shisheng Huang, Peihong Yu, Zhengxing Lian, Peixiang Ma, Jia Liu

## Abstract

Cytidine base editors (CBE) are novel genome engineering tools that can generate C-to-T nucleotide substitutions without introducing double-stranded breaks (DSBs). Instead of generating single-point mutations, CBEs induce nucleotide substitutions at wobble positions within the 20-nucleotide target site. A variety of strategies have been developed to improve the targeting scope and window of CBEs. Among these strategies, molecular switches that can temporally control CBE activities represent compelling options. In this study, we investigated the feasibility of using a bacteriophage-derived peptide, referred to as G8P_PD_, as the off-switch of CBEs. We showed that G8P_PD_ could be employed to control the activities of and improve the targeting window of A3A and BE3 CBEs and adenine base editor 7.10 (ABE7.10). Notably, in a cell-based disease model of Marfan syndrome, G8P_PD_ facilitated A3A CBE-based gene correction with a more focused targeting window and improved the percentage of perfectly edited gene alleles from less than 4% to more than 38% of the whole population. Our study presents the first peptide off-switch that can improve the targeting scope of CBEs, thus highlighting the importance of the temporal control of BE activity for precision base editing.

## Introduction

Clustered regularly interspaced short palindromic repeats (CRISPR)-CRISPR-associated genes (Cas) is the bacterial adaptive immune system for protecting host organisms from invading pathogens^1–3^. Owing to the modular feature, type II CRISPR systems, particularly CRISPR-Cas9, have been widely used for genome editing, transcriptional and epigenetic modulation and molecular imaging^4^. CRISPR-Cas9 can be directed to human genome by single-guide RNA (sgRNA) to create double-stranded breaks (DSBs) at targeted genomic loci^5–8^. In human cells, DSBs are repaired by two competing DNA repair pathways: error-prone nonhomologous end joining (NHEJ) and homology directed repair (HDR). The latter repair pathway can facilitate CRISPR-Cas9-mediated gene correction of pathogenic mutations^4^.

Recent studies have highlighted base editors (BEs) as an efficient genome editing tool for precision gene therapy^9–11^. BEs are fusion proteins comprising a catalytically inactive Cas nuclease and a nucleobase deaminase^12,13^. Unlike CRISPR-Cas9 genome editing tools, BEs generate precise base substitutions without introducing DSBs, thus avoiding concurrent, competing NHEJ events that incorporate nucleotide insertions and deletions (indels)^14^. Cytidine base editors (CBEs) and adenine base editors (ABEs) have been developed to realize genomic alterations of C-G to T-A and A-T to G-C, respectively. Besides DNA editing, Cas variants with RNA-binding capacities have been adapted for programmable RNA editing^15,16^.

The widely used third-generation CBEs, referred to as BE3, are fusion proteins composed of rat APOBEC (rAPOBEC), Cas9 D10A nickase and uracil glycosylase inhibitor (UGI). BE3 has a five nucleotide targeting window ranging from position 4 to 8, counting the protospacer adjacent motif (PAM) trinucleotides as positions 21 to 23^12^. An engineered CBE variant A3A, where rAPOBEC is replaced with human APOBEC3A (hAPOBEC3A), has a wider targeting window at positions 2 to 13 and is capable of editing methylated genomic regions^17^. Engineering endeavors have been made to improve the editing window of both BE3 (rAPOBEC-nCas9-UGI)^18^ and A3A (hAPOBEC3A-nCas9-UGI) CBEs^19^. Compared with ABEs, CBEs have higher genome-wide off-target events^20,21^. Despite of the considerable studies, the origin of the flexible editing windows and genome-wide off-target of CBEs are not completely understood.

It has been well known that excessive dosage of Cas9 and sgRNA can lead to elevated off-target mutations^22^. This finding led to the widespread use of directly delivered Cas9-guide RNA (gRNA) ribonucleoproteins (RNPs), which can improve the genome editing specificity by limiting the exposure of genome to gene-editing agents^23^. Similarly, delivery of BE-gRNA RNP can increase the base editing specificity compared with transfection of BE-coding plasmids^24^. Besides the use of gene-editing agents with short intracellular half time^23^, molecular switches provide compelling opportunities to improve the specificity of CRISPR-Cas via the temporal control of its cellular activity. Indeed, a variety of CRISPR-Cas inhibitors have been developed, including naturally occurring anti-CRISPR (Acr) proteins^25–27^, synthetic oligonucleotides^28^ and small molecules^29^. These inhibitors can improve the genome editing specificity of Cas9 RNPs^30^ and modulate the cellular activity of BEs^29^.

In a previous study, we have identified bacteriophage-derived peptides G8P_PD_ as inhibitors to *Streptococcus pyogenes* Cas9 (SpCas9)^31^. Herein, we hypothesized that the broad targeting window of CBEs is attributed, at least in part, to the excess intracellular dosage of base editing agents that could be alleviated by timed delivery of G8P_PD_. Indeed, we found that overexpression of the peptide inhibitors of SpCas9 in human cells could direct BE3 and A3A CBEs to more focused editing windows and facilitate precision gene correction in a cell model of Marfan syndrome^32^.

## Results

### G8PPD peptides inhibit CBE targeting of EGFP reporter

We have reported in a previous study that the periplasmic domain of the major coat proteins (G8P_PD_) from M13 and f1 bacteriophages (Fig. 1a) can inhibit the *in vitro* and cellular activities of SpCas9^31^. Because these peptides suppress SpCas9 activity by disrupting SpCas9-sgRNA binding^31^, we hypothesized that they could also act as inhibitors to cytidine base editors (CBEs) the activity of which also rely on sgRNA-guided DNA binding of SpCas9 domain. In order to evaluate the inhibitory activities of M13 and f1 G8P_PD_ on CBEs, we constructed an enhanced green fluorescent protein (EGFP) reporter carrying an inactivating mutation Y66C at the chromophore (Supplementary Fig. 1a). An sgRNA was designed to target the Y66C mutation site for introducing C-to-T conversion by CBE (Supplementary Fig. 1b). This mutation can correct EGFP-Y66C to wild-type genotype, yielding fluorescent cells that can be readily detected by flow cytometry. Inhibition of A3A CBE by G8P_PD_ peptides will lead to reduced EGFP fluorescence (Supplementary Fig. 1a).

**Fig. 1:**
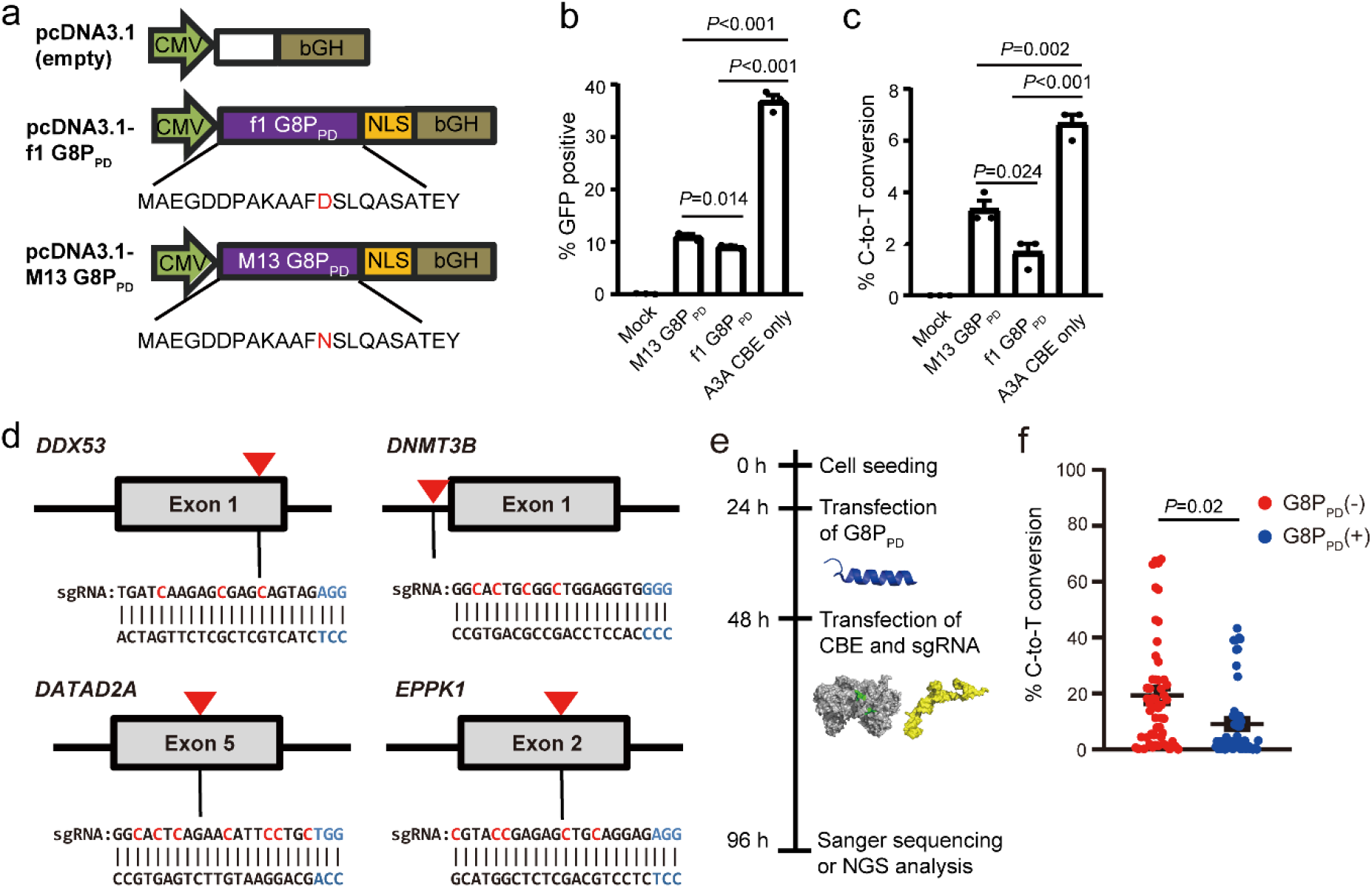
G8PPD peptides inhibit CBE targeting of EGFP reporter and endogenous genomic sites in human cells. **a** The design of G8P_PD_ plasmids for mammalian expression. **c** Flow chart showing experimental procedures. **b-c** The inhibitory effects of M13 and f1 G8P_PD_ toward A3A CBE in EGFP reporter cells, as determined by Sanger sequencing. **d** Design of sgRNA targeted to *DDX53, DNMT3B, GATAD2A* and *EPPK1* genomic loci. The PAM sequence and cytidines are highlighted in blue and red, respectively. Red arrows denote the positions of selected target sites. **e** Flow chart showing experimental procedures. The cartoons are blue for G8P_PD_, grey for CBE and yellow for sgRNA, respectively. **f** Analysis of the frequency of C-to-T conversion at all editing positions in the absence and presence of G8P_PD_. Significant difference between the groups with and without f1 G8P_PD_ is determined using two-tailed Student’s *t* test.

Our previous study has shown that G8P_PD_ peptides bind to sgRNA-free SpCas9 (apo-Cas9) and that maximum SpCas9-inhibiting activity of G8P_PD_ can be achieved by pre-incubating cells with overexpressed G8P_PD31_. Hence, in the present study we transfected G8P_PD_ plasmids at 24 h prior to the transfection of CBE^17^, sgRNA and EGFP-Y66C plasmids (Supplementary Fig. 1c). It was found that A3A CBE targeting of EGFP-Y66C reporter plasmid could recover the fluorescence in more than 30% HEK293T cells (Fig. 1b and Supplementary Fig. 1d). Pre-incubation of cells with M13 and f1 G8P_PD_ overexpression plasmid reduced fluorescent cells to approximately 10% of the whole population (Fig. 1b and Supplementary Fig. 1d), which confirmed our hypothesis that M13 and f1 G8P_PD_ could also function as inhibitors to BE. G8P_PD_-free pcDNA3.1 vector was transfected into HEK293T cells at 24 h prior to A3A CBE and sgRNA transfection to control for cell stress induced by serial transfections. Sanger sequencing of the PCR amplification product of A3A CBE-targeted site revealed evident C-to-T mutations in EGFP reporter (Fig. 1c and Supplementary Fig. 1e), consistently suggesting that M13 and f1 G8P_PD_ could inhibit CBE activity in human cells. It appeared that f1 G8P_PD_ has significantly higher inhibitory activity than M13 G8P_PD_ (Fig. 1b-c). Thus, subsequent studies are performed with f1 G8P_PD_.

### G8PPD inhibits CBE targeting at endogenous genomic sites in human cells

To examine the inhibitory effects of G8P_PD_ on CBE at endogenous sites, we designed sgRNA targeted to human *EPPK1*, *GATAD2A, DNMT3B* and *DDX53* genes respectively for A3A CBE (Fig. 1d). Similar to the procedure in EGFP reporter assay, f1 G8P_PD_-encoding plasmid was transfected into HEK293T cells at 24 h prior to A3A CBE and sgRNA transfection (Fig. 1e). The sgRNA-encoding and f1 G8P_PD_-encoding plasmids carried GFP and mCherry reporter genes respectively. In order to enrich cells with transfected plasmids, we used flow cytometry to enrich GFP and mCherry-dual positive cells (Supplementary Fig. 2) for analyses of CBE-induced genomic mutations.

Next-generation sequencing (NGS) analyses of sorted HEK293T cells revealed efficient base editing by A3A CBE at the four selected genomic sites (Supplementary Fig. 3a). The presence of f1 G8P_PD_ reduced CBE-mediated C-to-T conversion by various degree across different positions and genomic loci (Supplementary Fig. 3a). Statistical analyses of the editing efficiency at all examined positions revealed significant inhibitory activity of G8P_PD_ toward A3A CBE (Fig. 1f). Comparing different target sites or sgRNA, we found that the averaged inhibition rates of each target site had no significant difference (Supplementary Fig. 3b), suggesting that the CBE-inhibiting activity of G8P_PD_ was not dependent on genomic loci or sgRNA.

To investigate whether the activity of G8P_PD_ was cell-type dependent, we examined the performance of G8P_PD_ in U-2 OS cells. It was found that G8P_PD_ reduced the editing efficiency of A3A CBE by various degree across different genomic sites and editing positions (Supplementary Fig. 4a) in U-2 OS. Similar to the results in HEK293T cells, the inhibitory activity of G8P_PD_ in U-2 OS cells was not dependent on genomic loci or sgRNA (Supplementary Fig.4b). Collectively, these results suggested that G8P_PD_ could inhibit the editing activity of A3A CBE at different genomic sites across different cell types.

### G8PPD differentially inhibits the on-target and out-of-window editing of A3A CBE

We next analyzed the inhibitory activity of G8P_PD_ at different editing positions within each target site. It was found that the inhibitory effects of G8P_PD_ displayed notable variations along the 20-bp targeting site for each sgRNA. This result was consistently observed in HEK293T and U-2 OS cells (Fig. 2a). This observation prompted us to compare the effects of G8P_PD_ on the editing positions 4 to 8, which are conventionally deemed as the on-target editing window of CBEs, and on the out-of-window editing positions 1 to 3 and 9 to 20. It was found that G8P_PD_ significantly inhibited both the on-target and out-of-window editing of A3A CBE in HEK293T cells, with the latter being reduced to minimum level (Fig. 2b). Interestingly, G8P_PD_ had minor or little inhibition toward the on-target editing of A3A CBE but significantly inhibited the out-of-window editing (Fig. 2b).

**Fig. 2:**
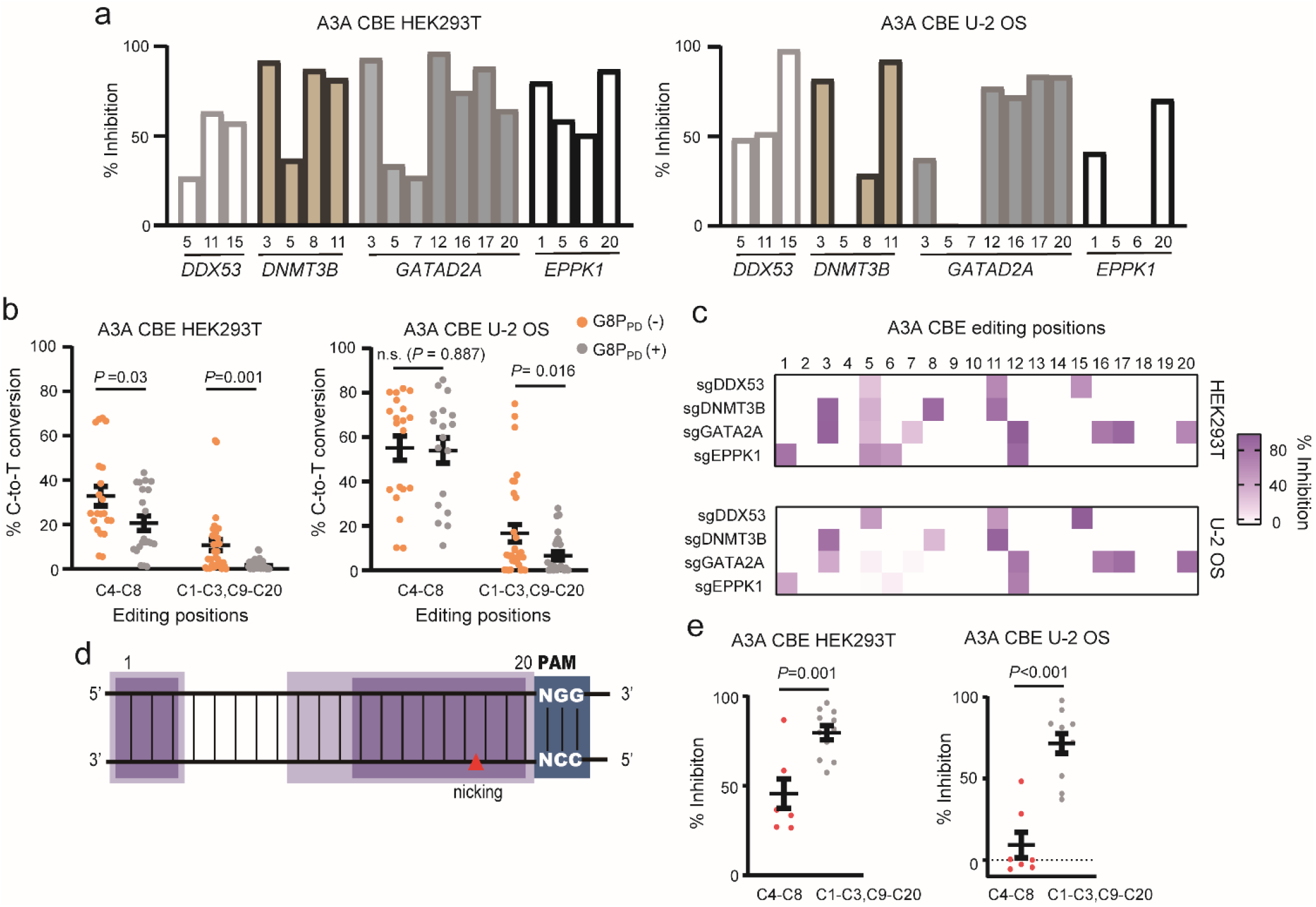
The inhibitory activities of f1 G8PPD toward A3A CBE at the endogenous genomic sites in human cell lines. **a** The inhibitory activity of f1 G8P_PD_ toward A3A CBE in HEK293T and U-2 OS cells. **b** Position-dependent inhibitory activity of G8P_PD_ at on-target and out-of-window editing positions of A3A CBE. Significant difference is determined using two-tailed Student’s *t* test. n.s., not significant. **c** Heat map showing position-dependent inhibition of A3A CBE by G8P_PD._ **d** Schematic presentation of the canonical out-of-window position (light purple) and the hot spots of G8P_PD_ inhibition (dark purple). PAM sequences are highlighted in blue. **e** Comparison of the inhibition rates of G8P_PD_ at the on-target and out-of-window editing positions of A3A CBE.

The differential effects of G8P_PD_ at the on-target and out-of-window editing positions were more evident when the inhibition rates were plotted along the 20-bp targeting site (Fig. 2c). Importantly, lowest inhibition rates were found at positions 4 to 8, the canonical on-target positions of CBEs (Fig. 2b). The hot spots of G8P_PD_-mediated inhibition had significant overlap with the canonical out-of-window editing positions of CBEs (Fig. 2d). Comparative analyses of the inhibition of C-to-T conversion showed that G8P_PD_ exhibited 2- and 10-fold selectivity of inhibition toward out-of-window positions over on-target positions in HEK293T and U-2 OS cells respectively (Fig. 2e).

### G8PPD differentially inhibits the on-target and out-of-window editing of BE3 CBE and ABE7.10

Because G8P_PD_ acts as a SpCas9 inhibitor, we envisioned that G8P_PD_ could inhibit different CBEs carrying SpCas9 module as the DNA-binding domain. Hence, we sought to examine the effects of G8P_PD_ on BE3 CBE and ABE7.10 that contain deaminase domains different from that in A3A CBE. We found that f1 G8P_PD_ could suppress BE3 CBE-induced C-to-T conversion though the inhibitory effects appeared to be less prominent than that with A3A CBE (Supplementary Fig. 5a). Similar to the results with A3A CBE, the inhibitory effects of f1 G8P_PD_ displayed varied inhibition rates at different editing positions of BE3 CBE (Fig. 3a), with the on-target positions 4 to 8 exhibiting lower inhibition compared to the out-of-window positions (Fig. 3b). It was observed that G8P_PD_ had 50-fold selectivity of inhibition toward the out-of-window positions over on-target positions (Fig. 3c).

**Fig. 3:**
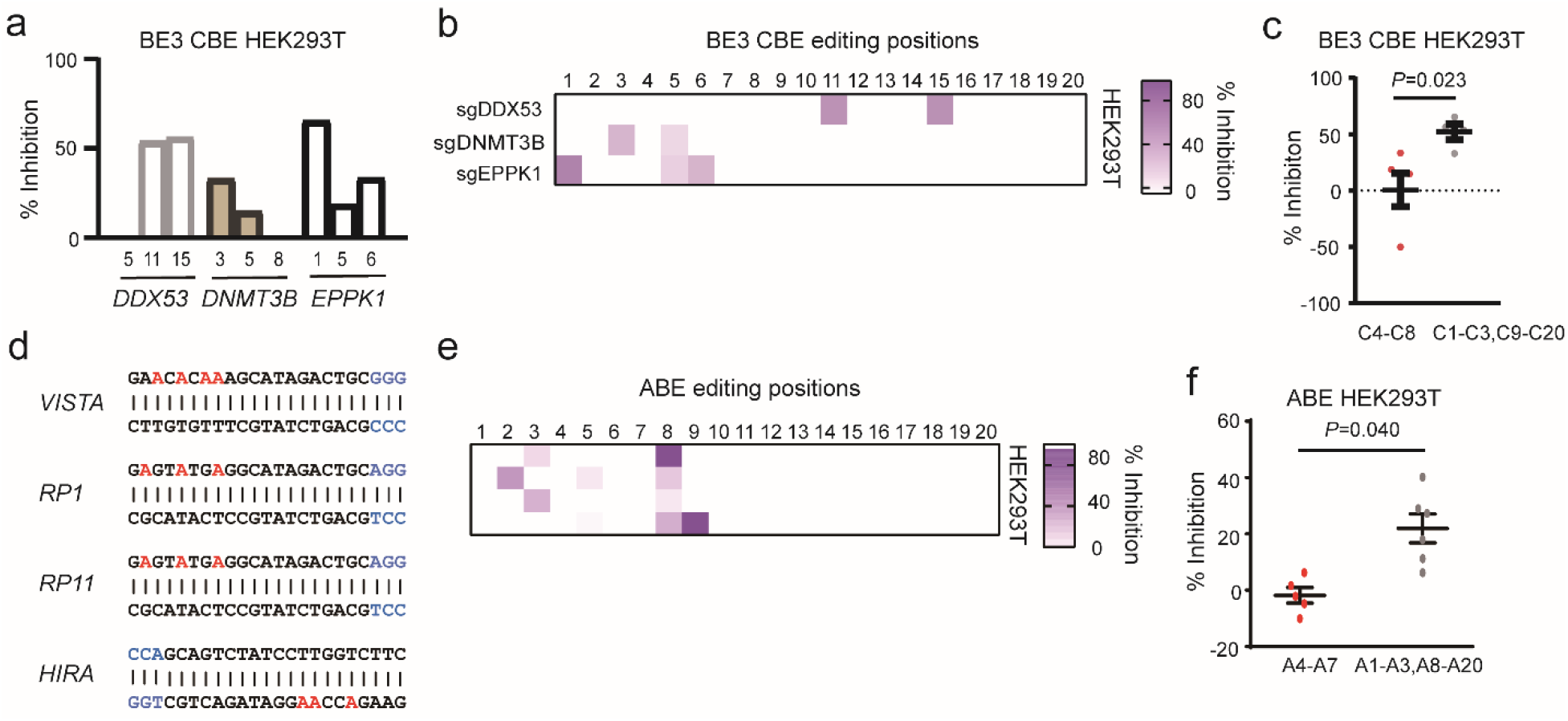
The inhibitory activities of f1 G8PPD toward BE3 CBE and ABE7.10 at the endogenous genomic sites in human cell lines. **a** The inhibitory activity of f1 G8P_PD_ toward BE3 CBE in HEK293T cells. **b** Heat map showing position-dependent inhibition of BE3 CBE by G8P_PD._ **c** Comparison of the inhibition rates of G8P_PD_ at the on-target and out-of-window editing positions of BE3 CBE. **d** Design of sgRNA for ABE7.10 targeted genomic loci. The PAM sequence and adenine positions are highlighted in blue and red, respectively. **e** Heat map showing the position-dependent inhibition of ABE7.10 by G8P_PD_. **f** Comparison of the inhibition rates of G8P_PD_ at the on-target and out-of-window editing positions of ABE7.10.

We next examined the effects of G8P_PD_ on ABE7.10. We designed four sgRNAs targeting to different genomic sites (Fig. 3d). Similar to the results with A3A and BE3 CBEs, f1 G8P_PD_ inhibited the A-to-G conversion activity of ABE7.10 in a position-dependent manner with positions 4 to 7, which is conventionally deemed as optimal on-target positions^13^, exhibiting lowest inhibition rates (Fig. 3e). f1 G8P_PD_ showed 20-fold selectivity of inhibition toward the out-of-window positions over on-target positions of ABE7.10 (Fig. 3f). These results collectively demonstrated that G8P_PD_ could preferentially inhibit the editing activities of CBE and ABE at out-of-window positions, suggesting the potential application of G8P_PD_ as an agent to improve the targeting scope of BEs.

As previous studies suggested that timed delivery of Acrs could improve the genome-editing specificity of SpCas9^30^, we next sought to investigate the effects of AcrIIA4 on modulating CBE and ABE activities. HEK293T cells were co-transfected with AcrIIA4 plasmid and sgRNA- and BE-coding plasmids (Supplementary Fig. 6a) and the inhibitory activity of AcrIIA4 was examined. It was found that AcrIIA4 suppressed the on-target and out-of-window activities of ABE and CBE to undetectable levels (Supplementary Fig. 6b) and that the strong inhibitory activities were observed across all positions within each sgRNA or genomic site (Supplementary Fig. 6c-d).

### G8PPD mediates precision correction of pathogenic FBN1 mutation by CBEs

To explore the potential therapeutic application of G8P_PD_, we assessed the effects of G8P_PD_ on the editing activities of A3A and BE3 CBEs in a cell-based disease model of Marfan syndrome^32^. In this cell model, a T7498C mutation was introduced into the *FBN1* gene of HEK293T cells to model the pathogenic amino acid mutation C2500R^32^. To enable CBE-mediated gene correction, we first designed sgRNA for A3A and BE3 CBEs to convert the pathogenic mutation T7498C to wild type (Fig. 4a). HEK293T-*FBN1*^T7498C^ cells were transfected with A3A and BE3 CBEs respectively in the absence or presence of G8P_PD_. Sanger sequencing of the PCR amplification products of *FBN1* gene from edited cells revealed efficient CBE-induced C-to-T conversion (Fig. 4b). NGS analysis showed that A3A and BE3 CBEs induced C-to-T conversion with varied frequencies across the 20-bp target site (Fig. 4c). Similar to the previous results, G8P_PD_ displayed remarkably higher inhibition rates at out-of-window positions than at on-target positions (Fig. 4d), with 36- and 3-fold selectivity for A3A and BE3 CBEs respectively (Fig. 4e). Most importantly, G8P_PD_ reduced the frequency and the number of genotypes of incorrectly edited alleles of A3A and BE3 CBEs (Supplementary Figs.7-8) and increased perfectly edited alleles of A3A CBE from less than 4% to more than 38% of the whole population, the latter of which corresponds to more than 50% of the total edited alleles (Fig. 4f). This 9-fold improvement demonstrated the feasibility of using G8P_PD_ as an agent for precision gene correction in Marfan syndrome model.

**Fig. 4:**
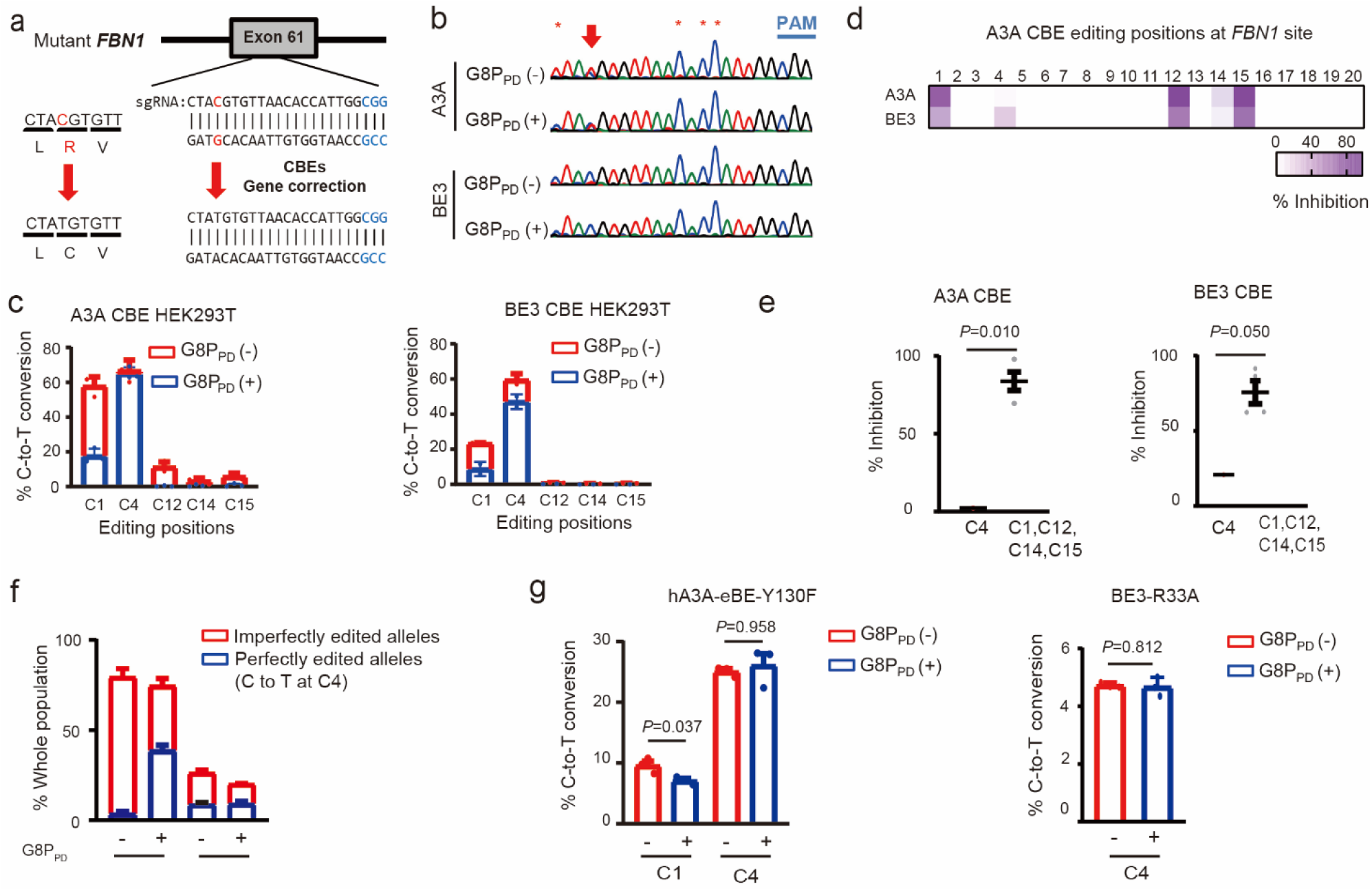
G8PPD improves the targeting window of CBEs for correction of *FBN1^T7498C^* mutation. **a** Schematic illustration of sgRNA design and the strategy for CBE-based gene correction. **b** Sanger sequencing of the positive strand of *FBN1* illustrating CBE-induced C-to-T mutations. Arrows and asterisks denote on-target and out-of-window cytidine positions for base editing. **c** Frequency of A3A and BE3 CBE-mediated C-to-T conversion in the absence and presence of f1 G8P_PD_. **d** Heat map showing the position-dependent inhibition of A3A and BE3 CBEs by G8P_PD_. **e** Comparison of the inhibition rates of G8P_PD_ at the on-target and out-of-window editing positions of A3A and BE3 CBEs at *FBN1* site. Significant difference is determined using Student’s *t* test. **f** Analyses of perfectly edited *FBN1* alleles in the absence and presence of G8P_PD_. Unless noted otherwise, the data are shown as mean ± SEM (*n* = 2 or 3) and significant difference is determined using two-tailed Student’s *t* test. **g** The effects of f1 G8P_PD_ on hA3A-eBE-Y130F and BE3-R33A CBEs for correction of *FBN1*^T7498C^ mutation.

We also investigated the effects of G8P_PD_ on high-fidelity CBE variants for correction of *FBN1*^T7498C^ mutation. It was found that G8P_PD_ could modestly inhibit the out-of-window editing activity of hA3A-eBE-Y130F at C1 position without affecting its on-targeting activity at C4 position (Fig. 4g). For BE3-R33A that exhibited only on-target editing at C4 position, G8P_PD_ did not significantly affect its activity. These results together suggested that peptide inhibitors of SpCas9 such as G8P_PD_ are compatible with and may even improve high-fidelity BE variants for precision gene correction.

## Discussion

It has well documented that CRISPR-Cas systems are associated with off-target cleavage^33^, chromosomal rearrangement^34^ and genotoxicity^35^. These adverse effects may result from uncontrolled expression of Cas proteins^22,36,37^. Unfortunately, most therapeutically relavant applications of CRISPR-Cas involve constitutively active Cas nucleases that raises major safety concern^38^. With the increasing number of therapeutic applications of CRISPR-Cas, the ability to manipulate the activity of CRISPR-Cas in human cells has become urgently needed. One feasible approach is to develop CRISPR-Cas off-switches to control the intracellular activity of Cas proteins. Thus far, synthetic oligonucleotides^28^, small-molecule inhibitors^29^ and bacteriophage-derived anti-CRISPR proteins (Acrs)^31^ have been developed to enable the temporal control of CRISPR-Cas activity. It has been shown that Acrs can improve the targeting specificity of SpCas9 at on-target sites over off-target sites^30^. This improvement was achieved via precisely controlled, timed delivery of Acrs along with Cas9-sgRNA ribonucleoproteins (RNPs)^30^. Yet, the effects of Acrs on base editors have not been explored. Notably, small-molecule inhibitors of SpCas9 have been employed as inhibitors to Cas9-based transcription factors and base editors^29^.

In our previous study, we discovered that the *in vitro* and cellular activity of SpCas9 can be inhibited by the periplasmic domain of bacteriophage major coat proteins (G8P_PD_)^31^. These G8P_PD_ peptides disrupt Cas9-sgRNA assembly by binding to the PAM interacting (PI) domain of SpCas9^31^. It was found that time-controlled delivery of overexpressed G8P_PD_ could improve the specificity of SpCas9 in human cells^31^. In the present study, we hypothesized that the out-of-window base editing of CBE may be attributed to excess CBE fusion proteins. Hence, we sought to explore whether timed delivery of SpCas9-inactivating G8P_PD_ could inhibit and improve the activities of CBE and ABE, in a rationale similar to that with the genome editing of SpCas9^31^. Our results have demonstrated that G8P_PD_ can preferentially inhibit the out-of-window editing of A3A and BE3 CBEs and ABE7.10, leading to more focused targeting window. To the best of our knowledge, our study represents the first peptide inhibitors that can improve the targeting scope of BEs. In comparison with G8P_PD_, AcrIIA4 abolished the activities of CBE and ABE at both on-target and out-of-window sites without any selectivity. These results support the notion that weak inhibitors, rather than strong inhibitors, are optimal agents for modulating the specificity of genome-editing tools. Nevertheless, it is possible that the editing activity of BEs may be improved by limiting the exposure time of SpCas9 to Acrs where Acrs are delivered at later time points during transfection.

It has been reported that the targeting specificity of CBEs and ABEs can be improved by incorporating genetic mutations into deaminase domain^20,21^. These BE variants are extremely useful when precision base editing is needed for therapeutic applications. In the present study, we have shown that G8P_PD_ has minor yet significant inhibition at the out-of-window editing position of hA3A-eBE-Y130F CBE. Most importantly, G8P_PD_ dose not affect the on-target editing activities of hA3A-eBE-Y130F and BE3-R33A CBEs. These results have established the feasibility of G8P_PD_ not only as an alternative approach to improving the targeting specificity of BEs but also as an additive agent to combine with BE variants. In further studies, it would be interesting to evaluate the inhibitory activities of G8P_PD_ toward dual adenine and cytidine base editors (A&C-BEs)^39^ and prime editors^40^. These studies can advance our understanding of the capability of G8P_PD_ serving as general modulators of CRISPR-Cas genome engineering tools.

Although we only examined f1 and M13 G8P_PD_ for inhibition of CBE activity in the current study, there might be other G8P_PD_ from inoviridae phages^31^ that can function as CBE off-switches. Comparative analyses can be performed on these different G8P_PD_ peptides to determine the sequence-activity relationship. This information would facilitate the design and development of next-generation peptide off-switches of CBEs. Moreover, in-depth understanding of the mechanism of action of G8P_PD_ on CBE inhibition can also advance their therapeutic applications as modulators for precision gene correction.

## Methods

### Plasmid construction

sgRNA was cloned into pGL3-sgRNA expression vector carrying a U6 promoter and an EGFP reporter gene (Addgene, #107721). CBE-targeted genomic sites are indicated in Supplementary Table 1. The sequence of sgRNA-encoding oligonucleotides were listed in Supplementary Table 2. Human codon-optimized DNA sequences encoding M13 and f1 G8P_PD_ were cloned into the *BamH* I/*Xba* I sites of pcDNA3.1(+) by plasmid recombination kit Clone Express (Vazyme Biotech, Nanjing, China). These G8P_PD_ peptides carry a C-terminal SV40 nuclear localization signal (NLS) for co-localization with Cas9 proteins. G8P_PD_ peptides were cloned into plv-EF1α-mCherry plasmid harboring mCherry fluorescent protein marker. For construction of EGFP-Y66C reporter plasmid, Y66C mutation was introduced by quickchange PCR.

### Cell culture and transfection

HEK293T and U-2 OS cells were obtained from ATCC and maintained in Dulbecco's Modified Eagle Medium (DMEM) (Cat. No. SH30243, Hyclone, Logan, USA) supplemented with 10% fetal bovine serum (FBS) and 1% penicillin-streptomycin at 37 °C with 5% CO_2_. Transfection was performed using lipofectamine 2000 Reagent (Cat. No. 11668019, ThermoFisher Scientific, Waltham, USA) according to manufacturer’s instructions. HEK293T and U-2 OS cells were seeded on to poly-D-lysine (Cat. No. A-003-E, Sigma, St. Louis, USA) pre-coated 24-well plates at 24 h prior to transfection.

For EGFP reporter assay, 1 μg G8P_PD_-encoding plasmid was transfected into HEK293T cells. After 24 h, 0.25 μg sgRNA plasmid, 0.5 μg A3A CBE plasmid and 0.05 μg the EGFP-Y66C reporter plasmid were co-transfected into G8P_PD_-expressing cells. At 48 h after the second transfection, cells were harvested and analyzed by flow cytometry. For base editing at endogenous genes, cells of approximately 70% confluency were transfected with 1 μg G8P_PD_ plasmid that carries an mCherry selection marker at 12 h after seeding. At 24 h after G8P_PD_ transfection, 0.25 μg sgRNA plasmid that carries a GFP selection marker and 0.5 μg CBE plasmid were transfected in G8P_PD_-expressing cells. At 48 h after CBE and sgRNA plasmids transfection, cells were harvested and analyzed by Beckman Coulter CytFLEX (Beckman Coulter, Brea, USA) or sorted by BD FACSAria III flow cytometry (BD Biosciences, New York, USA). At least 2,000 mCherry and GFP dual positive cells were collected for subsequent analyses. Analyses of the Sanger sequencing results of the PCR product of edited EGFP reporter were performed using EditR^41^.

For AcrIIA4 co-transfection experiments, HEK293T cells were transfected with 0.5 μg of CBE plasmid and 0.25 μg sgRNA-expressing and 1 μg of AcrIIA4-expressing plasmids that carry GFP and mCherry reporters, respectively. At 72 h after transfection, mCherry and GFP dual positive cells were collected for subsequent analyses.

### Extraction and PCR amplification of genomic DNA

Genomic DNA was extracted using QuickExtract™ DNA Extraction Solution (Lucigen, USA). Genomic PCR was performed using 100 ng genomic DNA, corresponding primers (Supplementary Table 3-4), Phanta Max Super-fidelity DNA Polymerase (Cat. No. P505-d1, Vazyme) or KOD plus (Cat. No. F0934K, Takara, Kyoto, Japan) with a touchdown cycling protocol that contains 30 cycles of 98 °C for 10 s, X °C for 15 s where X decreases from 68 °C to 58 °C with a −1 °C/cycle rate and 68 °C for 60 s.

### NGS analyses of PCR amplicons

PCR was performed using the NGS primers (Supplementary Table 4). PCR products were purified using Gel Extraction Kit (Cat. No. D2500-02, OMEGA) before construction of NGS libraries. Hiseq3000 SBS&Cluster high-throughput NGS library preparation kit (Cat. No. FC-410-1002, Illumina, San Diego, CA, USA), (VAHTS Universal DNA Library Prep Kit (ND608-01, Vazyme) or TruSeq NanoDNALT Library Prep Kit (Illumina) were used to generate dual-indexed sequence following the manufacturer’s protocol. Briefly, more than 50 ng purified PCR fragment was used for direct library preparation. The fragments were treated with End Prep Enzyme Mix (Cat. No. WE0229, Illumina) for end repairing, 5′ phosphorylation and dA-tailing in one reaction, followed by a T-A ligation to add adaptors to both ends. Size selection of adaptor-ligated DNA was then performed using VAHTSTM DNA Clean beads (Cat. No. N411-03, Vazyme) or Beckman AMPure XP beads (Cat. No. A63882, Illumina). Each sample was then amplified by PCR for 8 cycles using P5 and P7 primers (Supplementary Table 4). Both P5 and P7 primers carry the sequences that can anneal with flow cells for bridge PCR. In addition, P7 primer carries a six-base index allowing for multiplexing. The PCR products were cleaned using VAHTSTM DNA Clean beads (Cat. No. N411-03, Vazyme) or Beckman AMPure XP beads (Cat. No. A63882, Illumina), validated using an Agilent 2100 Bioanalyzer (Agilent Technologies, Palo Alto, CA, USA) and quantified by Qubit2.0 Fluorometer (Invitrogen, Carlsbad, CA, USA) or Quant-iT PicoGreen dsDNA Assay Kit (ThermoFisher). Two or three biological replicates were processed by Genewiz (Suzhou, China) or Personalbio (Shanghai, China) using Illumina HiSeq 3000. Sequencing was carried out using a 2 × 150 paired-end (PE) or 2 × 300 paired-end (PE) configuration. Image analyses and base calling were conducted by the HiSeq Control Software (HCS) + OLB + GAPipeline-1.6 (Illumina) on the HiSeq instrument. Sequencing reads were obtained in the Fastq format. Amplicons with less than 6 M read counts were excluded from the analyses.

To obtain the editing efficiencies, the adapter pair of the pair-end reads were removed using AdapterRemoval version 2.2.2, and pair end read alignments of 11 bp or more bases were combined into a single consensus read. All processed reads were then mapped to the target sequences using the BWA-MEM algorithm (BWA v0.7.16). For each site, the mutation rate was calculated using bam-readcount with parameters −q 20 −b 30. Indels were calculated based on reads containing at least 1 inserted or deleted nucleotide in protospacer. Indel frequency was calculated as the number of indel-containing reads/total mapped reads.

## Data availability

High-throughput sequencing data has been deposited in the NCBI Sequence Read Archive database under the accession code PRJNA601083. Additional materials or information can be obtained from J.L. upon reasonable request.

## Statistical analyses

For cell-based assay, three biological replicates are generally performed and the results are shown as mean ± standard error of the mean (SEM). For NGS analyses, two or three biological replicates are performed. Statistical analyses were performed using two-tailed Student’s *t* test unless otherwise noted.

## Acknowledgements

We thank the High-Throughput Screening Platform and Biomedical Big Data Platform at Shanghai Institute for Advanced Immunochemical Studies (SIAIS) at ShanghaiTech University for the support of flow cytometry experiments and analyses of NGS data. This work is supported by the National Natural Science Foundation of China (31600686 to J.L.) and ShanghaiTech University Startup Fund (2019F0301-000-01 to J.L.)

## Author contributions

J.L. conceptualized study. J.L., K.J. and Y.-R.C. designed the experiments, S.H. analyzed next-generation sequencing data. K.J. and Y.-R.C. performed the *in vivo* CBE-inhibiting experiments. Z.L. and P.M. provided critical resources. J.L. and K.J. and Y.-R.C. wrote the manuscript. All authors discussed the results and approved the manuscript.

